# Comparison of High-Throughput Single-Cell RNA Sequencing Data Processing Pipelines

**DOI:** 10.1101/2020.02.09.940221

**Authors:** Mingxuan Gao, Mingyi Ling, Xinwei Tang, Shun Wang, Xu Xiao, Ying Qiao, Wenxian Yang, Rongshan Yu

## Abstract

With the development of single-cell RNA sequencing (scRNA-seq) technology, it has become possible to perform large-scale transcript profiling for tens of thousands of cells in a single experiment. Many analysis pipelines have been developed for data generated from different high-throughput scRNA-seq platforms, bringing a new challenge to users to choose a proper workflow that is efficient, robust and reliable for a specific sequencing platform. Moreover, as the amount of public scRNA-seq data has increased rapidly, integrated analysis of scRNA-seq data from different sources has become increasingly popular. However, it remains unclear whether such integrated analysis would be biased if the data were processed by different upstream pipelines. In this study, we encapsulated seven existing high-throughput scRNA-seq data processing pipelines with Nextflow, a general integrative workflow management framework, and evaluated their performances in terms of running time, computational resource consumption, and data processing consistency using nine public datasets generated from five different high-throughput scRNA-seq platforms. Our work provides a useful guideline for the selection of scRNA-seq data processing pipelines based on their performances on different real datasets. In addition, these guidelines can serve as a performance evaluation framework for future developments in high-throughput scRNA-seq data processing.

## 1 Introduction

Since the emergence of the first single-cell RNA sequencing (scRNA-seq) platform [1], many research achievements have been made at the cellular and subcellular levels with unprecedented resolutions with the aid of this technology. Recent advances in microfluidics and next generation sequencing (NGS) have further increased the efficiency and throughput of scRNA-seq, enabling more cells to be identified and the expression information of more genes for each cell to be quantified simultaneously [2, 3, 4]. From micro-capillary pipettes in 2012 [5, 6] and nanoliter droplet-based microfluidic chips in 2015 [7, 8] to the latest liquid barcoding combined with split-pool methods [9], the number of cells analysed in parallel has increased from several to tens of thousands [10], bringing new challenges for the processing of a large amount of barcoded NGS data for efficient and accurate quantification of the transcript information of single cells.

To meet the needs of high-throughput scRNA-seq data processing, several pipelines that integrate multiple functions have been developed. As one of the first high-throughput scRNA-seq platforms, Drop-seq [7] was introduced in 2015. Moreover, Drop-seq-tools in combination with Picard tools were introduced and provided a user-friendly Application Programming Interface (API) to process the data from Drop-seq. Later in 2017, three additional pipelines were published including Cell Ranger [11], umis [12] and UMI-tools [13]. Cell Ranger [11] was developed along with the widely adopted 10X Genomics platform which can process multiple biological samples separated by sample barcodes in parallel. For fair comparison of different scRNA-seq platforms, umis [12] was developed as a versatile pipeline to analyse data produced by over 15 sequencing platforms. Using this uniform pipeline, accuracy, sensitivity and other critical metrics were calculated using the External RNA Controls Consortium (ERCC) samples to establish a performance benchmark of the scRNA-seq platforms. UMI-tools [13] systematically modelled the UMI errors occurring in scRNA-seq data and used a directional network UMI correction strategy to achieve more accurate quantification results both on simulated and real data. More recently, three flexible and comprehensive pipelines were published, namely, dropEst [14], scPipe [15] and zUMIs [16, 17]. DropEst was specifically designed to process data acquired from high-throughput droplet-based scRNA-seq platforms. ScPipe implements the upstream data processing pipeline as a Bioconductor R package, with additional APIs for subsequent downstream analysis. ZUMIs integrates almost all available data formats with or without UMI and can be applied to data from 21 different sequencing platforms (version 2.4.5b). Moreover, this all-in-one pipeline considers intron-mapping reads to improve transcript counting.

The progression of various high-throughput scRNA-seq data processing pipelines has empowered end-users, including biologists, chemists and developers of novel scRNA-seq platforms, to process their data conveniently and challenged them to select a proper pipeline for their data in hand. Recently, several studies have been published to evaluate scRNA-seq data analysis strategies [18, 19, 20, 21, 22, 23, 24, 25, 26]. These studies mainly focused on downstream analysis algorithms including normalization, data imputation, clustering, and differential expression. For upstream analysis, namely, analysis performed by data processing pipelines, the performance of three different genome aligners was investigated [27]. However, comparison of the performance of whole scRNA-seq data processing pipelines that integrate multiple functions and procedures on real datasets remains to be conducted.

Here, we focused on the evaluation of seven existing high-throughput scRNA-seq data processing pipelines. To facilitate our study, we encapsulated all seven scRNA-seq data processing pipelines using Nextflow [28], which provided not only a user-friendly tool for large-scale project management but also an integrated package for rapid deployment and convenient utilization of different pipelines. After that, we compared the performances of these pipelines in terms of their computational efficiency and consistency of their biological analyses on various real datasets. The results of our study could serve as a valuable reference for researchers to select proper pipelines for their applications. The software and example data used in this study are made publicly available at https://github.com/xmuyulab/scRNAseq_pipelines to further enable users to perform benchmarking studies on their own data.

## 2 Methods

### 2.1 Datasets and software

#### 2.1.1 Datasets

The following datasets were used in our experiments.

- **Drop-HM dataset**: Drop-seq 50 cells per microliter human (HEK) and mouse (3T3) mixed dataset (GEO: GSE63269)
- **Drop-ERCC dataset**: Drop-seq ERCC dataset (GEO: GSE66694)
- **Seq-Well-PBMC dataset**: Seq-Well PBMC dataset (GEO: GSE92495)
- **Quartz-SVF dataset**: Quartz-seq2 mouse stromal vascular fraction (SVF) dataset (GEO: GSE99866)
- **inDrop-ERCC dataset**: inDrop v1 ERCC dataset (GEO: GSE65525)
- **10X-PBMC-10k dataset**: 10X Genomics v3 10k peripheral blood mononuclear cell (PBMC) from a healthy donor (https://support.10xgenomics.com)
- **10X-PBMC-5k dataset**: 10X Genomics v3 5k peripheral blood mononuclear cell (PBMC) from a healthy donor (https://support.10xgenomics.com)
- **10X-HM dataset**: 10X Genomics v3 1k 1:1 human (HEK) and mouse (3T3) mixture (https://support.10xgenomics.com)
- **10X-ERCC dataset**: 10X Genomics 1k ERCC dataset (https://support.10xgenomics.com)

#### 2.1.2 Software packages evaluated

The versions and source urls of the seven pipelines that we evaluated are listed below.

- **Drop-seq-tools** version-2.3.0 (https://github.com/broadinstitute/Drop-seq/releases)
- **Cell Ranger** version-3.0.2 (https://github.com/10XGenomics/cellranger)
- **scPipe** version-1.4.1 (https://github.com/LuyiTian/scPipe)
- **zUMIs** version-2.4.5b (https://github.com/sdparekh/zUMIs)
- **UMI-tools** version-1.0.0 (https://github.com/CGATOxford/UMI-tools)
- **umis** version-1.0.3 (https://github.com/vals/umis)
- **dropEst** version-0.8.6 (https://github.com/hms-dbmi/dropEst)

#### 2.1.3 Auxiliary software tools used in experiments

In addition, we used the following data processing and alignment tools.

- **Picard tools** version-2.18.14 (http://broadinstitute.github.io/picard)
- **STAR** version-2.7.0f (https://github.com/alexdobin/STAR) [29]
- **Rapmap** version-0.6.0 (https://github.com/COMBINE-lab/RapMap) [30]
- **SAMtools** version-1.9 (https://github.com/samtools/samtools) [31]
- **Nextflow** version-19.04.1 (https://www.nextflow.io)

### 2.2 Overview of data processing pipelines

High-throughput scRNA-seq data processing pipelines typically implement multiple steps, including sequencing data demultiplexing, alignment and transcript quantification, to transform raw NGS data in BCL or fastq file formats to expression matrices for further downstream analysis [32] (Figure 1). In a high-throughput scRNA-seq experiment, transcripts from tens of thousands of cells can be collected in a single sequencing library. Transcripts from different cells are then converted to complement DNA (cDNA) via reverse transcription and labelled with cell barcodes and UMIs [33, 34, 35] for the isolation of cells and molecules, respectively. In the processing pipelines, de-multiplexing is first performed on raw reads from the sequencer to extract their cell barcodes to retrieve single cell information, as illustrated in Figure 1. The processing of cell barcodes and UMIs affects not only the accuracy of transcript quantification of each cell but also the compatibility of the data processing pipeline on different scRNA-seq platforms.

**Figure 1:**
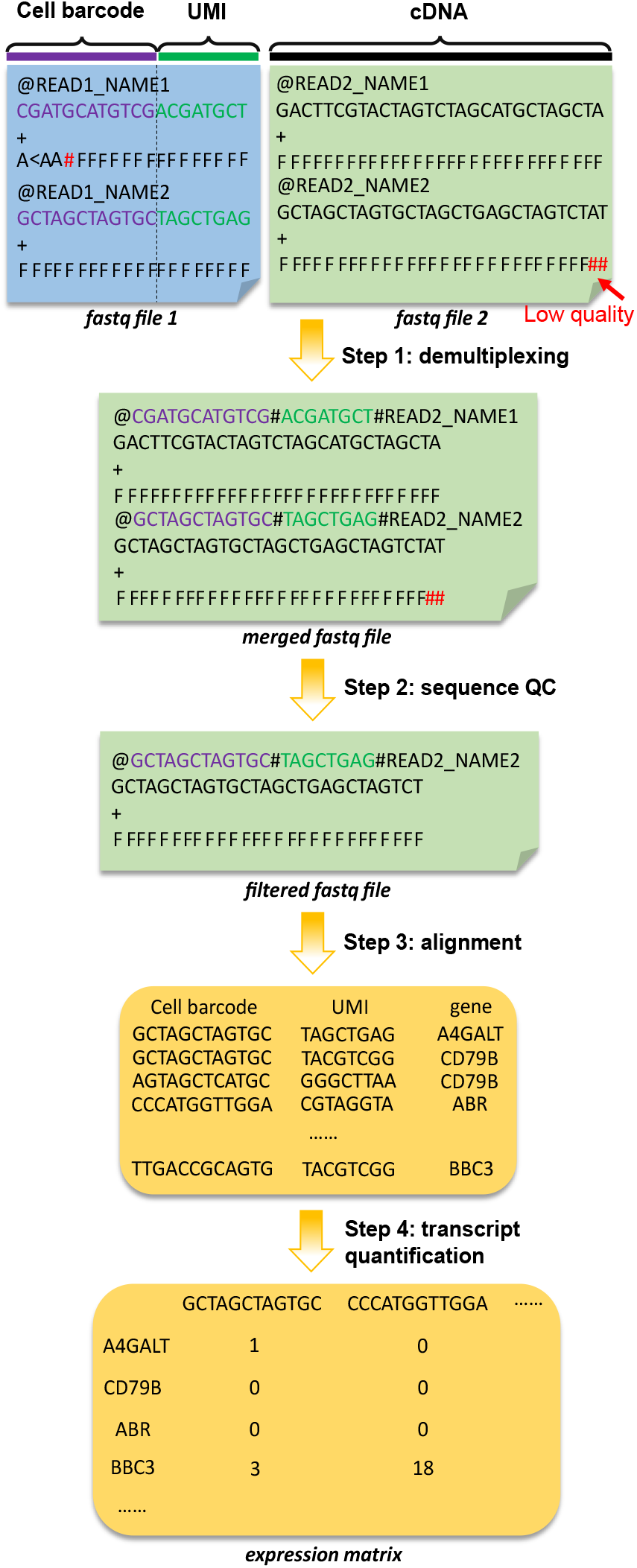
The major steps of high-throughput scRNA-seq data processing pipelines, namely, demultiplexing, sequence QC, alignment, and transcript quantification, that are applied to generate an expression matrix representing the gene expression of single cells from sequencing reads.

After demultiplexing, quality control (QC) was performed on the raw reads. Pipelines except umis and dropEst directly discard reads with one or more low-quality bases of cell barcodes and UMIs to avoid adverse effects from sequencing error, since cell barcodes and UMIs contain essential read information and cannot be trimmed as cDNA sequences. For umis and dropEst, only reads with low-quality cell barcodes are discarded. Moreover, Drop-seq-tools suggests trimming adapters and other interfering sequences, such as poly-A of cDNA reads, while the other pipelines do not perform quality-based cDNA pretrimming by default.

The processed reads are further aligned to a reference genome or transcriptome. Many aligners developed for bulk RNA-seq can also be applied to scRNA-seq data [36]. Umis utilizes pseudo-alignment tools such as Kallisto [37] and Rapmap [30]. ScPipe uses Subread [38] by default. The other pipelines included in this study use STAR [29] as a default option for alignment. Furthermore, some pipelines, such as UMI-tools, zUMIs and scPipe, are compatible with multiple aligners.

Finally, aligned reads are assigned to their corresponding cell barcodes, and the expression matrices can be constructed by counting the unique reads with the same cell barcode, UMI and gene. PCR duplicates, which are indicated by reads with identical cell barcodes and UMIs, are not counted. To address possible barcode synthesis errors and sequencing errors, different error handling and correction strategies are used in different pipelines. For Drop-seq-tools, umis, zUMIs and scPipe, cell numbers can be decided with prior knowledge of the experimental design, heuristic elbow plots of cumulative read counts of each cell or predefined whitelists of cell barcodes. Cell barcodes with an inadequate number of reads or not on the whitelist will be discarded directly without any correction. Other pipelines choose more aggressive approaches. UMI-tools do not correct any cell barcodes by default, while users can add the `--error-correct-cell` parameter to merge cell barcodes within a 1-Hamming distance. Cell Ranger considers cell barcodes that are within a 1-Hamming distance to predesigned cell barcodes based on the posterior probability calculated from base qualities of observed cell barcodes. For dropEst, UMI-gene composition similarity is modelled by Poisson distribution to determine whether cell barcodes within a 2-Hamming distance should be merged, and this method outperforms simply filtering cell barcodes with whitelists or the number of reads by analysing Drop-seq and 10X data.

Similarly, different UMI collapsing strategies have been adopted in different pipelines. Drop-seq-tools, zUMIs and scPipe collapse UMIs based on their Hamming distances, with cut-off values that can be specified by users. Cell Ranger corrects UMI errors based on the base quality of UMIs. UMI-tools systematically models UMI errors both on simulated data and real data to verify that the directional graph-based collapsing algorithm behaves better than other methods. DropEst designs a Bayesian approach for UMI error correction, where the posterior probability of error occurrence is estimated from factors including the prior UMI distribution, base quality, and gene expression.

A detailed comparison of these pipelines is included in Supplementary Table S1. Note that as different pipelines have different processing steps, we decided not to compare the performance of each individual step of different pipelines in this review. Instead, we will only focus on the overall performances of pipelines that are relevant to scRNA-seq studies.

### 2.3 Nextflow pipeline design and features

Nextflow [28] is an integrative workflow management framework using Apache Groovy script language. It has been widely used to build bioinformatics data analysis workflow with flexibility, extensibility and reproducibility [39, 40, 41, 42, 43, 44]. Herein, we used Nextflow to encapsulate the seven data processing pipelines so that users can directly call different pipelines through a uniform set of APIs. In addition, the developed Nextflow script also allows users to customize parameters or assign computing resources to individual steps of the pipelines. The entire workflow is further packed as a docker image with an Anaconda virtual environment so that it can be readily deployed to different platforms without laborious software compiling and running environment setups. All of the encapsulated pipelines are available online.

### 2.4 Data processing

#### 2.4.1 Evaluation of computational performance

We used the 10X-PBMC-10k dataset, which contains 640 million reads in total, to evaluate the computational performances of the pipelines. To test the performances under different input file sizes, we randomly sampled the dataset to construct reduced databases of different sizes from 0.5 million to 100 million reads. In our experiments, Rapmap was used in umis, while STAR was used in the other pipelines for alignment. The cell number was set to 10000 for the pipelines without automatic cell number determination algorithms (Supplementary Table S1). All the other parameters of the pipelines were set to their default values. All testing programs were run on Ubuntu 16.04 with a CPU of 64 cores and 256 Gb of memory. We made all CPU resources available on the testing machine during the test. Program execution information was collected from the Nextflow tracing and visualization API.

#### 2.4.2 Evaluation of transcript quantification performance

ERCC datasets from three different scRNA-seq platforms (Drop-ERCC, inDrop-ERCC and 10X-ERCC) were used to evaluate the transcript quantification performance of the pipelines. For the Drop-ERCC and 10X-ERCC datasets, we quantified the gene expression of 84 and 1015 common cells produced from all compatible pipelines. For the inDrop-ERCC dataset, since different pipelines (dropEst, umis and UMI-tools) produced different cell barcodes as the exact length of the cell barcodes in inDrop v1 data was unspecified [8], it was difficult to identify the common cells from them. Therefore, for this dataset, we compared the gene expression of 900 cells with the top UMI counts from the respective pipelines. To compare the gene expression levels, a linear regression model was built between the logarithmic expression counts and the logarithmic actual RNA-molecule concentrations (ground truthed) for each cell. R^2^ values were used to evaluate the transcript quantification accuracy of the pipelines.

We further tested these pipelines on two human and mouse mixed datasets (Drop-HM and 10X-HM) to evaluate their demultiplexing and transcript quantification performances. For the 10X-HM dataset, all seven pipelines were tested. For the Drop-HM dataset, however, six pipelines, not including Cell Ranger, were tested due to compatibility issues. Herein, we aligned the reads to the hg19 and mm10 mixed reference genome. After the expression matrices were generated, we discarded cells that were either multiplets, empty cells or noise. We classified cells with more than 80000 transcripts as multiplets and those with fewer than 5000 transcripts as empty cells or noise for the Drop-HM dataset, and the corresponding cut-off values for the 10X-HM dataset were set to 140000 and 2500 transcripts, respectively. Cells with species-specific transcript rates higher than 95% were annotated as human cells or mouse cells and mixed cells or vice versa. Other parameters passed to the pipelines were set to default values.

#### 2.4.3 Downstream analysis with Seurat

We used Seurat [45] for downstream analysis, including data normalization, feature selection, clustering, dimension reduction, and differential expression [46]. For each dataset, expression matrices produced by different pipelines were built into Seurat objects. Subsequently, cells were filtered according to the distributions of transcript counts and mitochondrial gene counts. For fairness, we only extracted common cells identified by all pipelines for subsequent analysis. Then, logarithmic standardization was performed with a constant scale factor (`scale.factor=10000`). Genes with high variation were selected by the FindVariableFeatures function (`nfeatures=2000`), and principal component analysis (PCA) was applied to these 2000 genes to extract 20 principal components (PCs). Subsequently, unsupervised clustering and nonlinear dimension reduction were performed on the 20 PCs of each cell. Cell identities were manually annotated by canonical markers for the resulting clusters. Differentially expressed genes were found by the FindAll-Markers function (`min.pct=0.25, logfc.threshold=0.25`) for each annotated cell type. Then, genes with adjusted p-values (Bonferroni correction) less than 0.05 were retained for further comparison. Imputation algorithms were not used in the analyses here to avoid potential bias introduced by other factors.

#### 2.4.4 Supervised cell type identification with SuperCT

In addition to manual cell type identification, we also used the results from SuperCT [47], a neural network-based supervised learning model, to evaluate the performances of different pipelines. In our experiment, we simply fed the expression matrices produced by all the pipelines to the SuperCT web application (https://sct.lifegen.com/) to identify the cell types automatically. Then, pairwise confusion matrices were used to evaluate the consistency of cell types identified by the different pipelines.

## 3 Results and discussion

### 3.1 Computational performance

We evaluated the computational performance of the seven pipelines in terms of running time, CPU usage and memory consumption. As shown in Figure 2A, the running time increases roughly linearly with the number of reads being processed in all pipelines except for umis, which has a significant constant overhead in the running time for small datasets. Among the seven pipelines, dropEst and scPipe showed the highest efficiency for small datasets with less than 100 million reads. For larger datasets, Cell Ranger and scPipe had the highest efficiency. umis was the slowest pipeline for small datasets due to its significant overhead. Further investigation of the individual steps of umis (Supplementary Figure S1) shows that its transcript quantification step accounts for most of the computational time and was irrelevant to the number of reads being processed. For large datasets, Drop-seq-tools was the slowest, requiring more than 24 hours to complete analysis of a dataset with 640 million reads.

**Figure 2:**
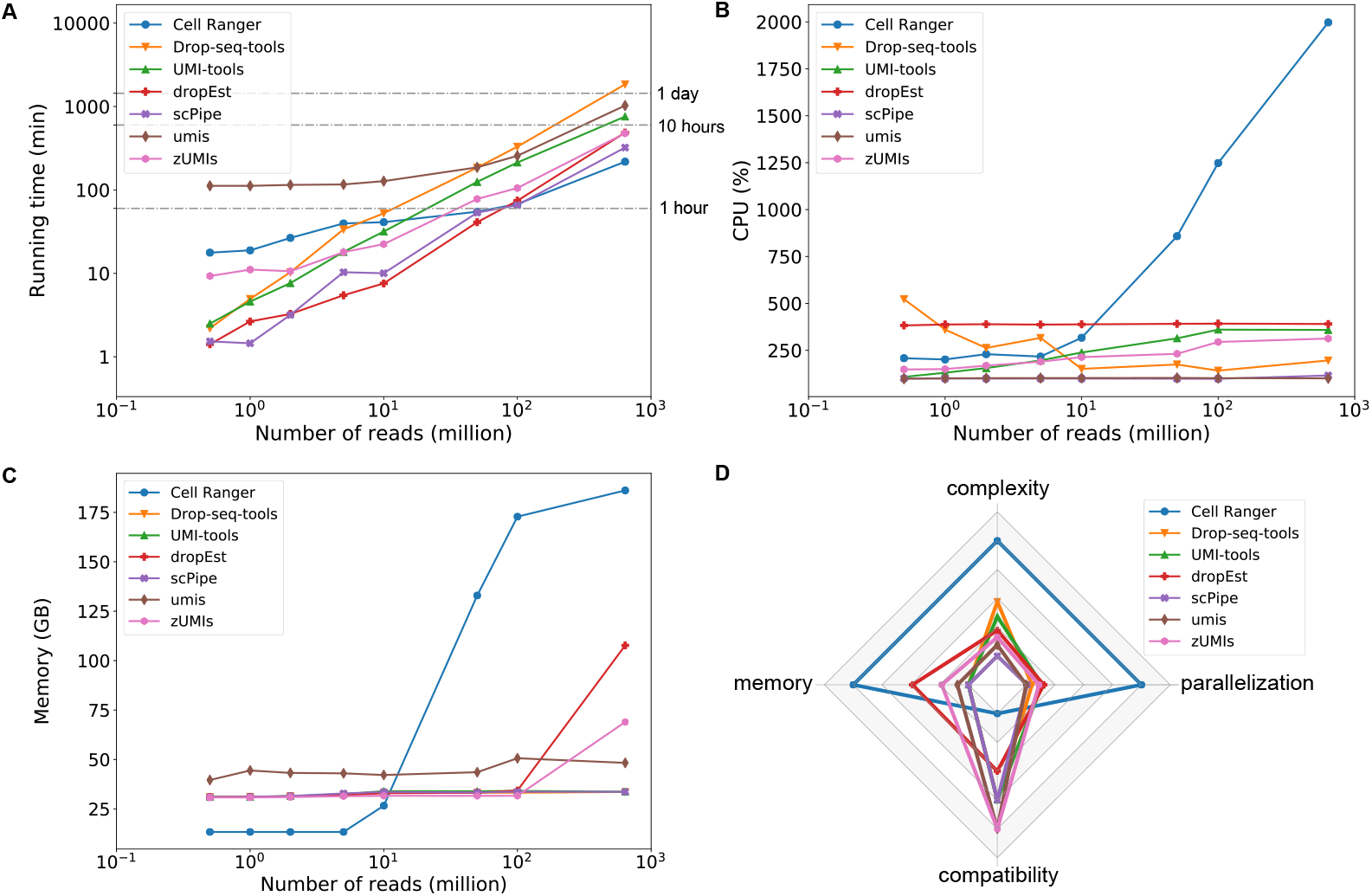
Comparison of the computational performances of the pipelines. (**A**) Running time without alignment steps, (**B**) maximum CPU usage, (**C**) maximum memory consumption, and (**D**) radar chart of the summary of the performances of different pipelines on a full 10X-PBMC-10k dataset with 640 million reads, where parallelization stands for the maximum of the CPU cores used, and algorithm complexity corresponds to running time×parallelization. The compatibility is indicated by four discrete levels including a single platform, droplet-based platforms, UMI-based platforms and almost all platforms (Supplementary Table S1).

Figure 2B compares the maximum CPU usage of the pipelines. As both Rapmap and STAR allow for manual setting of the number of CPU cores, we compared only the CPU usage of the remaining steps. Note that the peak CPU usage of Cell Ranger, zUMIs and UMI-tools increases with increasing amounts of reads. Among the pipelines evaluated, Cell Ranger demonstrated the highest level of parallelization when dealing with large datasets.

The maximum memory consumption of each pipeline was also investigated. As shown in Figure 2C, Cell Ranger required less memory than the other pipelines when processing small datasets with less than 10 million reads, while its memory usage drastically increased when the amount of data increased. The other pipelines showed relatively stable memory consumption, except for dropEst and zUMIs, which required more memory for processing datasets of 640 million reads.

Figure 2D provides a summary of three aspects of computational performance and compatibility for all pipelines based on their performance on a large dataset (640 million reads). Among them, Cell Ranger had the highest level of parallelization and algorithmic complexity, which make it favourable for high-speed processing of a large amount of data at the cost of high CPU and memory usages. scPipe, umis and zUMIs demonstrated lower algorithm complexity, which makes them favourable for large-scale scRNA-seq data analyses when computational resources are limited. In terms of platform compatibility, Cell Ranger only supports 10X data, while scPipe, umis and zUMIs are compatible with a wider range of platforms and hence are more suitable for large-scale integration studies.

### 3.2 Demultiplexing and transcript quantification

For high-throughput scRNA-seq, transcript quantification-based downstream analysis is currently the primary application due to the 3’-bias in library preparation [36, 48]. To evaluate the transcript quantification performances of different pipelines, we first used the ERCC datasets from three high-throughput scRNA-seq platforms. The ERCC standard spike-in mixture consists of 92 different RNA species with various predesigned lengths and concentrations [49], and has been used to evaluate bulk RNA-seq protocols [50, 51] as well as scRNA-seq platforms [12, 52]. Herein, we compared the linear relationship between the quantification results from different pipelines and input RNA-molecule concentrations of individual cells. As shown in Figure 3, the results of all pipelines were highly consistent for the 10X-ERCC dataset. For the other two datasets, differences between pipelines were more obvious, and UMI-tools achieved the best performance on both datasets with the highest mean accuracy and the lowest deviation among individual cells.

**Figure 3:**
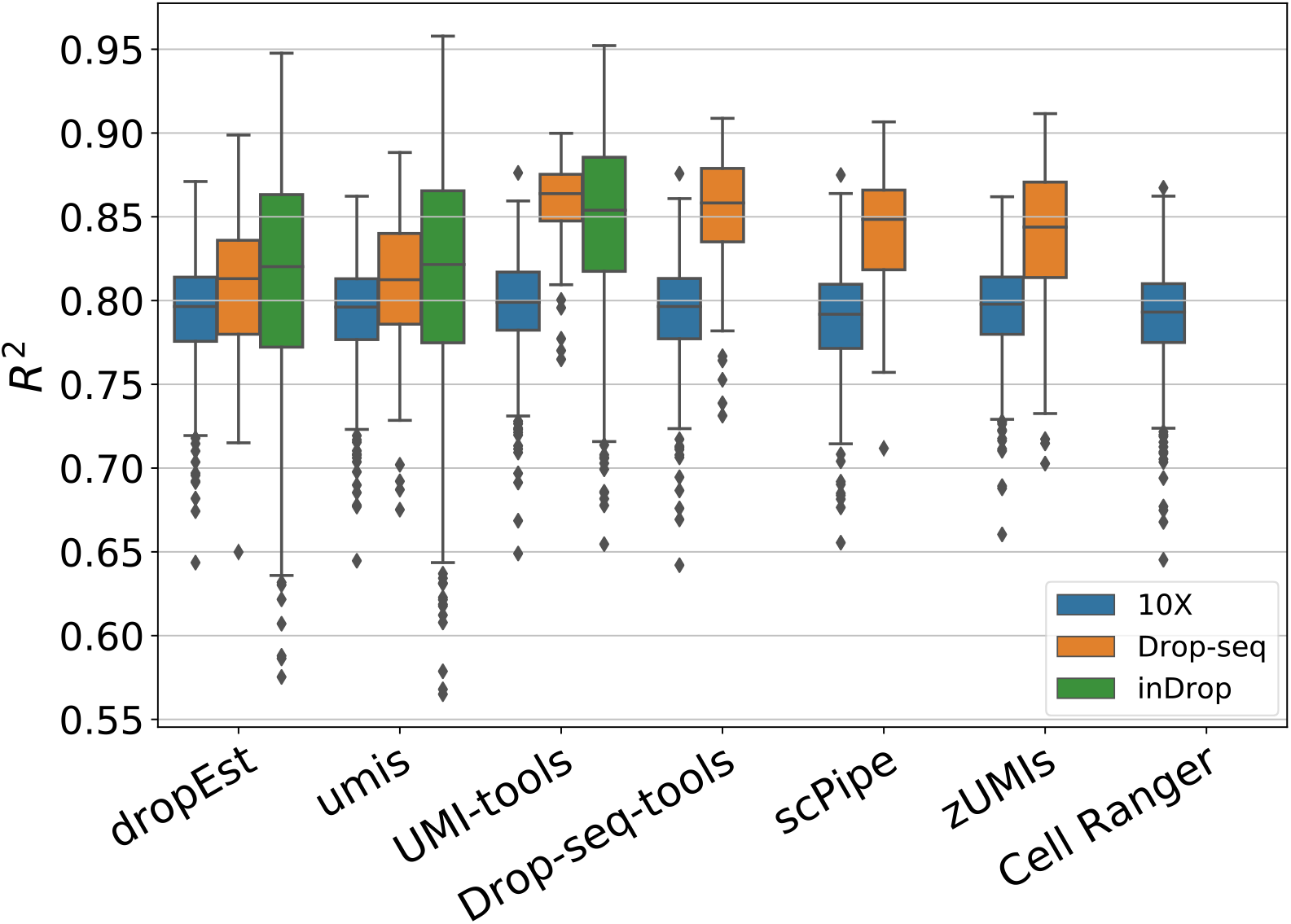
R^2^ values of the linear relationship between the quantification results from different pipelines and the input RNA-molecule concentrations. Note that some pipelines could not process the Drop-ERCC and/or inDrop-ERCC datasets due to compatibility issues.

We further evaluated the pipelines with a human and mouse mixed dataset (Drop-HM dataset) created by mixing a human cell line (HEK) and a mouse cell line (3T3), which is commonly used in evaluating the single cell isolation fidelity of high-throughput scRNA-seq platforms [7, 53, 54, 55]. Ideally, the quantification result of a single cell should only contain transcripts from either species, but not both. However, due to errors introduced in either sequencing and/or scRNA-seq data processing, it is possible that erroneous cells that contain transcripts of both species are present in the final expression matrices. First, the results show that the majority of the cells (953 out of 1350) generated by these pipelines were common, demonstrating high consistency in cell recovery among the six pipelines (Supplementary Figure S2A). Pairwise Pearson correlations of gene expression of the 953 common cells further showed that the results from Drop-seq-tools, UMI-tools and scPipe were highly similar (Figure 4), which was expected because these three pipelines share a very similar data processing strategy including aligner used and exon-only mapping and no cell barcode correction. Moreover, cells from umis showed different gene expression distributions compared with those from other pipelines. Supplementary Figure S3A shows that most cells from Drop-seq-tools and UMI-tools were located on either the X or Y axis, indicating that the major cells from these two pipelines were free of transcript contamination. The quantification results of scPipe also shared a similar pattern to those of Drop-seq-tools and UMI-tools, with slightly more cells with mixed transcripts at the regions of low transcript counts. In contrast, more off-axis cells were observed in the results for zUMIs and dropEst, suggesting that these two pipelines may introduce more transcript contamination than the pipelines mentioned above. Cells with relatively higher cross-species transcript contamination can also be observed in the results from umis, but to a lesser extent.

**Figure 4:**
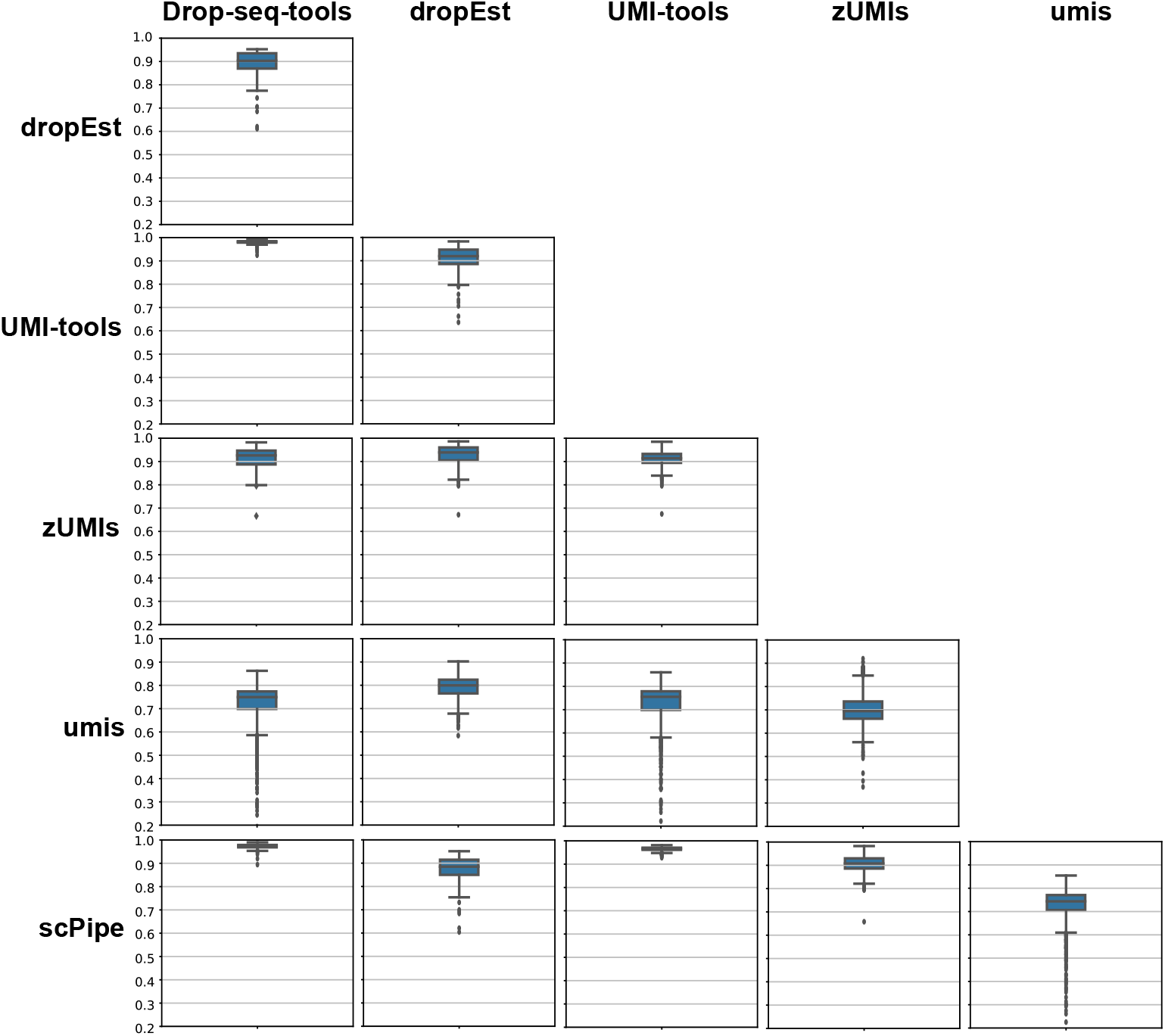
Pairwise Pearson correlation boxplots of the gene expression of 953 shared cells across seven pipelines for the Drop-HM dataset.

To further investigate the cross-species contamination effects of zUMIs and dropEst, we extracted mixed cells with more than 30000 mouse transcripts and fewer than 10000 human transcripts, resulting in 17 and 15 cells from these two pipelines, respectively. We then compared their transcript counts with those of the corresponding cells from Drop-seq-tools. The results show that both zUMIs and dropEst produced higher human and mouse transcript counts than Drop-seq-tools (Supplementary Figure S3B and Figure S3C), suggesting higher sensitivity of these two pipelines in transcript quantification. This may be explained by their relatively aggressive cell barcode and UMI correction algorithms. Note that the high sensitivity of these two pipelines may introduce adverse effects, such as cross-species transcript contamination, as illustrated in the mixed species experiment above.

We also performed the same experiment on the 10X-HM dataset to include Cell Ranger in the comparison, which was designed specifically for 10X data. The results showed that 924 cells out of 992 were shared by the seven pipelines (Supplementary Figure S2B). The cell barcode purity and Pearson correlation of different pipelines on the 10X-HM dataset are shown in Supplementary Figures S4 and S5, respectively. We observed that most pipelines produce relatively similar results compared to those from the Drop-HM dataset, which could possibly be due to the lower noise level of the pre-designed barcode of the 10X platform. However, the results from zUMIs and umis showed gene expression patterns that were slightly but not significantly different from those of the other pipelines.

Finally, the number of genes of single cells detected by different pipelines was compared with the Drop-HM dataset and 10X-HM dataset. ZUMIs and dropEst demonstrated higher sensitivity in transcript quantification by detecting more genes than the other pipelines, whereas umis detected the lowest number of genes in both datasets (Supplementary Figure S6).

### 3.3 Clustering and cell type identification

High-throughput scRNA-seq provides expression information for thousands of genes from a large number of cells per test, and it has long been challenging to identify cell types with such high dimensional data. The widely adopted approaches for cell type identification are mainly based on unsupervised clustering to assign cells into distinct groups, and the cell type of each group is further identified using canonical markers or differential genes by manual annotation [56, 57, 58]. Supervised learning algorithms have also been proposed to improve the efficiency and accuracy of cell type identification [24, 47, 59, 60, 61, 62, 63]. To better understand the effect of scRNA-seq data processing pipelines on cell type identification, we compared the performance of the unsupervised clustering-based method and SuperCT [47], a supervised learning-based method, using the expression matrices generated by different pipelines.

First, we combined the expression matrices generated by the seven pipelines for the 10X-PBMC-10k dataset and performed t-distributed stochastic neighbour embedding (tSNE) on the combined matrix. Here, we considered only cells common to all seven matrices, which gave us 9520 cells for each pipeline in total. The tSNE plots are given in Figure 5, in which cells are coloured by pipeline in Figure 5A and by cell type identified with SuperCT [47] in Figure 5B. We can see that the cells were clustered into distinct groups by pipeline rather than by cell type, except that cells from Drop-seq-tools and Cell Ranger were grouped together. The same experiment was also performed on the Seq-Well-PBMC dataset [54]. For this dataset, cells from Drop-seq-tools, scPipe, dropEst and UMI-tools were grouped together; differences among cells of different types can roughly be observed, but the boundaries were not clear. Cells from umis and zUMIs were clustered into distinct groups (Supplementary Figure S7). The results indicate that different pipelines can introduce additional unwanted confounding factors akin to the batch effect [64, 65, 66], which need to be considered when integrating expression matrices from different studies to avoid introducing noise that may mask true biological variation or create false positive results if the disturbances introduced by different processing pipelines are directional.

**Figure 5:**
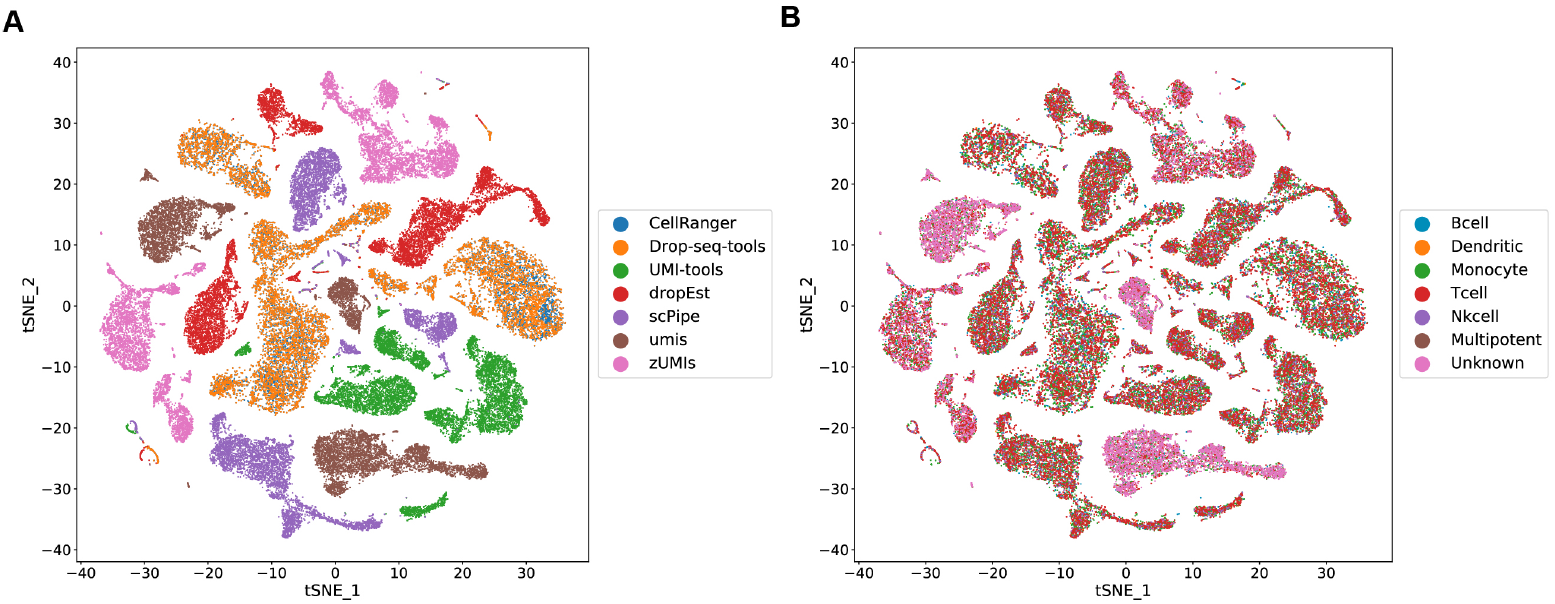
tSNE plots of the same cells from the 10X-PBMC-10k dataset identified by different pipelines (**A**) coloured by pipelines, and (**B**) coloured by cell types identified with SuperCT.

Next, we compared the cell types identified by SuperCT using the expression matrices from different pipelines. Cell types identified from Drop-seq-tools, UMI-tools, dropEst, Cell Ranger and scPipe were highly consistent for the 10X-PBMC-10k dataset, while the expression matrices from zUMIs and umis resulted in more “unknown” cells, which could possibly be due to differences in the distributions of expression matrices generated by zU-MIs and umis compared to the data used to train the supervised learning model (Figure 6 and Supplementary Figure S8). Therefore, more attention should be paid to choosing data processing pipelines when supervised learning based cell type identification models are going to be used in downstream analysis. A similar phenomenon can also be observed in the results from the Seq-Well-PBMC dataset (Supplementary Figure S9 and Figure S10).

**Figure 6:**
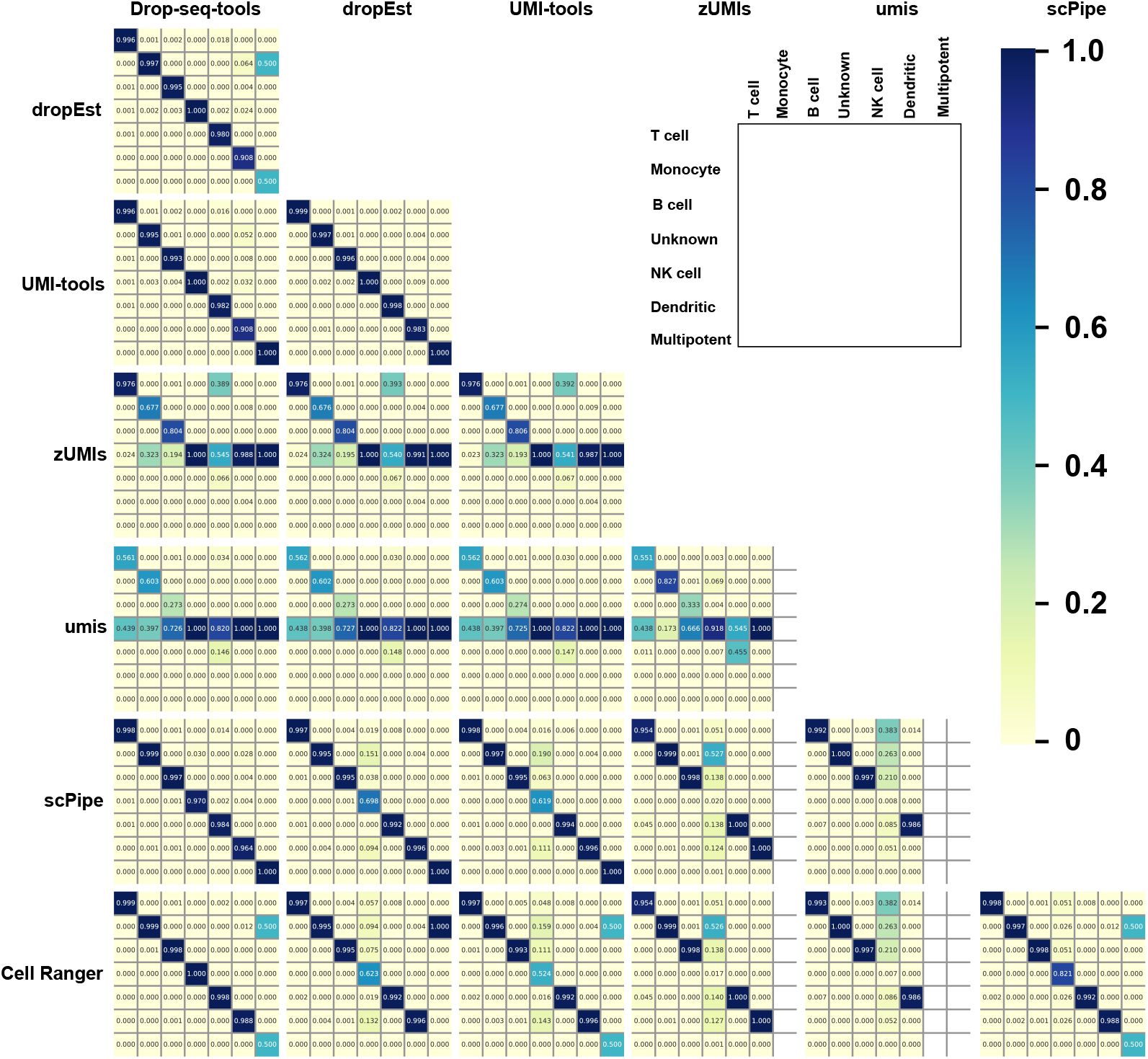
Confusion matrices of the cell types of the 10X-PBMC-10k dataset identified with SuperCT for seven pipelines.

Furthermore, we conducted the same test using the widely adopted unsupervised cell type identification method (Methods). Here, cell identities were manually annotated based on canonical markers by three different researchers independently. We only focused on major cell types with clearly defined canonical marker genes. Majority voting was used to resolve inconsistent annotations and reduce randomness brought by manual cell type annotation. As shown in Supplementary Figures S11 and S12, cell types annotated from different pipelines showed higher consistency on the 10X-PBMC-10k dataset and Seq-Well-PBMC dataset than with the supervised learning method. Among all the pipelines, the results from Drop-seq-tools, scPipe and UMI-tools showed the highest level of consistency, which was similar to the results of SuperCT above. Another dataset (Quartz-SVF dataset) that contained cell subtypes that SuperCT was not able to identify was also tested, and similar results were observed (Supplementary Figure S13).

### 3.4 Differential expression

Another important step of high-throughput scRNA-seq data analysis is to find differentially expressed genes (DEGs) across various cell populations [18]. Here, we compared the DEGs identified by Seurat using expression matrices from different pipelines on 10X-PBMC-10k and Seq-Well-PBMC datasets. The cell types were annotated by SuperCT in advance, and majority voting was used if a cell was annotated to different cell types by different pipelines. Only cells existing in all pipelines were considered.

As shown in Figure 7 and Supplementary Figure S14, the DEGs of each cell type from different pipelines were highly consistent, with those from zUMIs and umis tended to have slightly more unique genes than the other pipelines. We further checked the average logarithmic fold changes of several commonly used canonical markers [58, 67, 68] of each cell type for the different pipelines, which showed that zUMIs had lower average logarithmic fold changes for most signature genes than the other pipelines (Figure 8).

**Figure 7:**
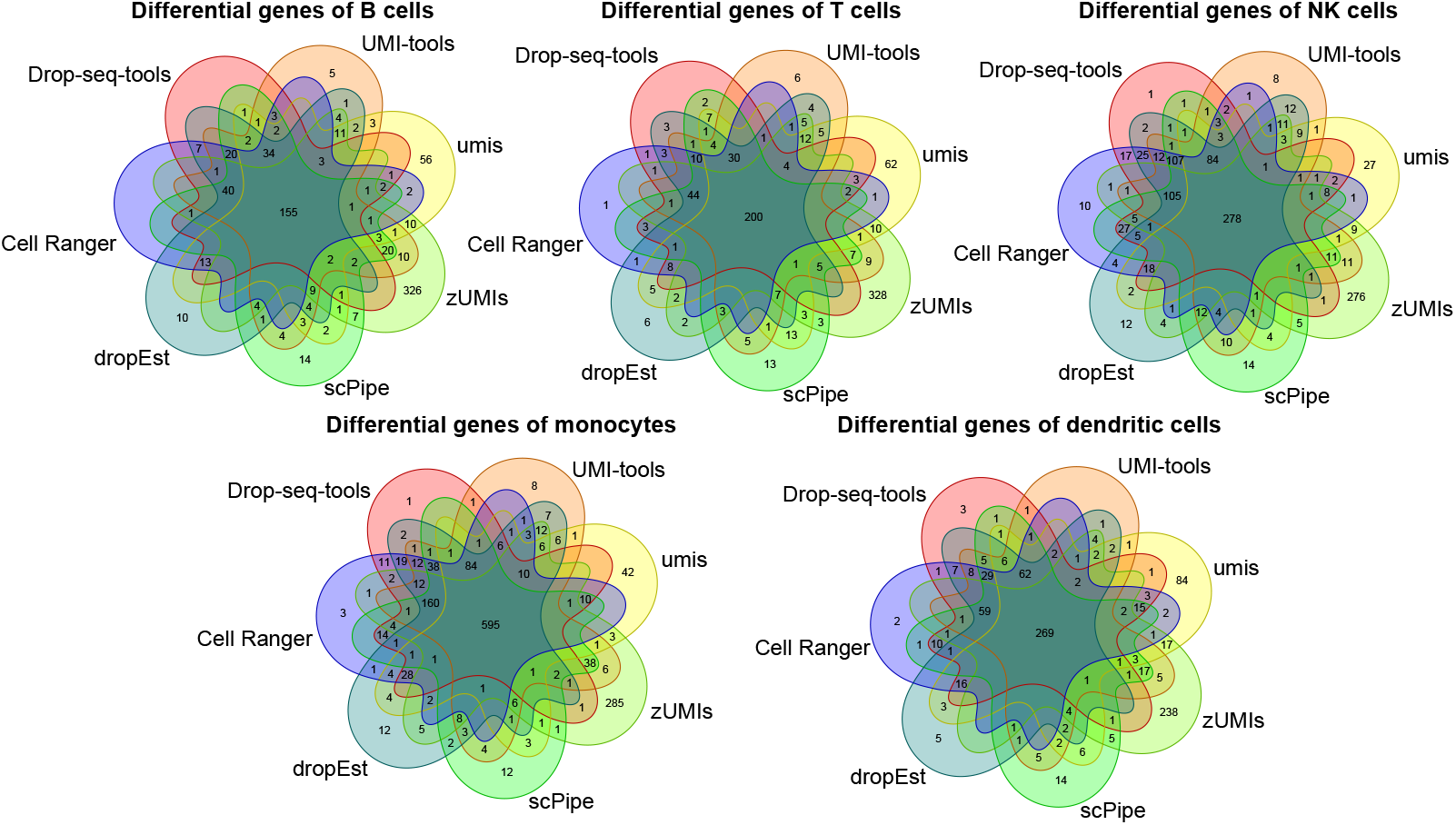
Venn diagrams of differentially expressed genes found by Seurat with an adjusted p-value (Bonferroni correction) less than 0.05 in the 10X-PBMC-10k dataset.

**Figure 8:**
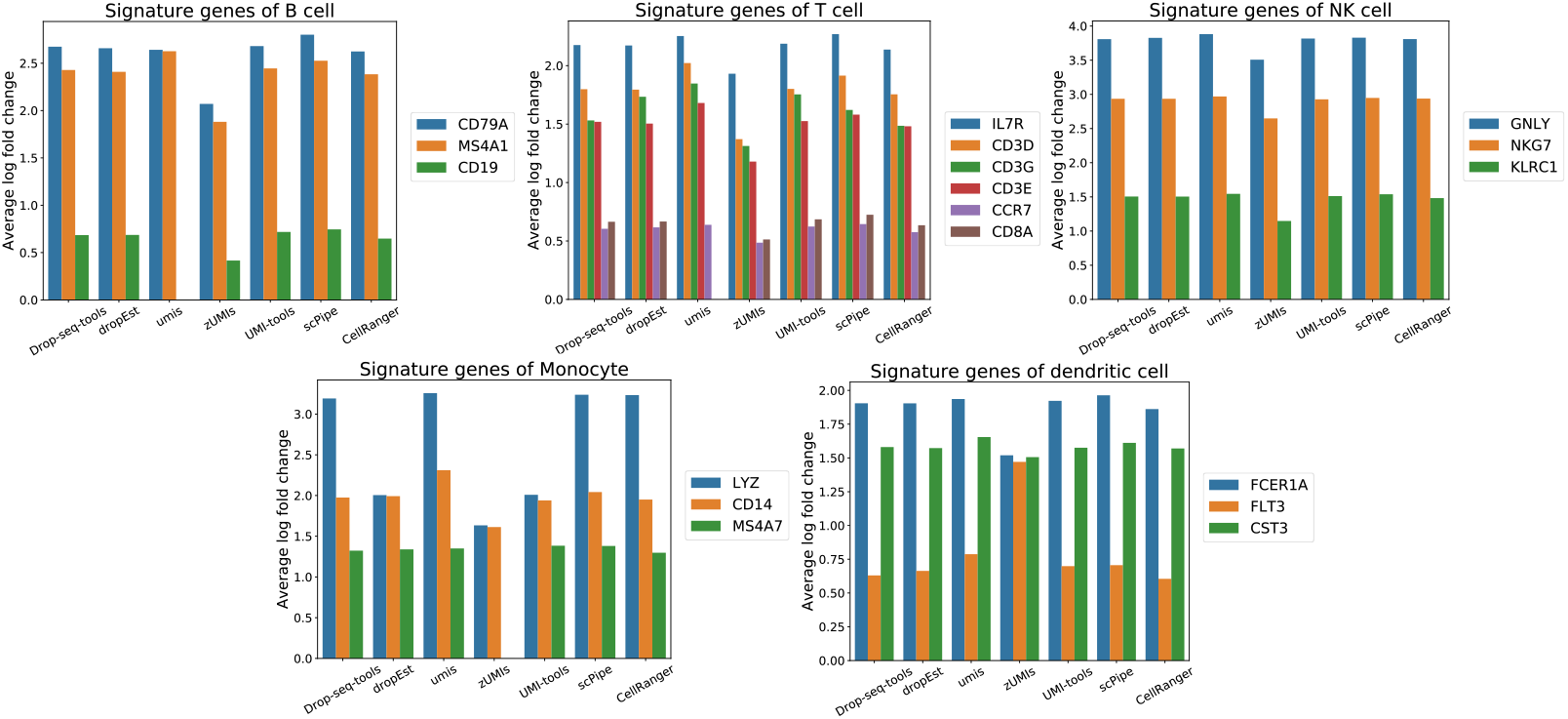
Average log fold changes of signature genes of different cell types from the 10X-PBMC-10k dataset. If a gene was not contained in the differential gene list, the average log fold change was set to zero.

## 4 Conclusions

This study reports a comprehensive review of seven high-throughput scRNA-seq data processing pipelines in terms of their computational and biological analysis performances using nine real datasets from five high-throughput scRNA-seq platforms.

First, for computational performance, Cell Ranger demonstrated the highest algorithm complexity if we count the total running time multiplied by the number of CPU cores. However, as Cell Ranger also has the highest level of algorithm parallelization, it is still able to achieve reasonable running time performance compared to other pipelines if a high-performance computer is available. On the other hand, scPipe, umis and zUMIs demonstrated lower algorithm complexity and good compatibility for different scRNA-seq platforms, suggesting their usability for large-scale scRNA-seq integration analysis when computational resources are limited. Moreover, Drop-seq-tools, scPipe and UMI-tools had the lowest levels of memory consumption.

Second, transcript quantification results from different pipelines were tested on ERCC spike-in datasets and human/mouse mixed datasets. For ERCC datasets, UMI-tools achieved the highest accuracy of transcript quantification for the data from three different scRNA-seq platforms. For human/mouse mixed datasets, all of the pipelines were able to correctly provide cells with low cross-contamination. ZUMIs and dropEst had higher sensitivities compared to the other pipelines as they tended to detect more genes and produce higher transcript counts for each cell. However, such high sensitivity in transcript quantification may bring unwanted confounding factors for subsequent analyses, as suggested in our human/mouse mixture experiment, where these two pipelines produced more erroneous cells with cross-species transcript contamination compared to other pipelines. Therefore, care should be taken to better balance sensitivity and specificity in judging quantification results from different analysis pipelines.

Third, the influence of pipelines on biological analysis was tested on cell type identification and differential expression analysis. Our results show that expression matrices generated by different pipelines cannot be directly integrated for downstream analysis due to the variation caused by the pipelines. For both supervised learning-based models and unsupervised clusetering-based cell type identification methods, the results from Drop-seq-tools, scPipe and UMI-tools show high consistency compared with those from the other pipelines. Specifically, zUMIs and umis showed relatively different distributions of identified cell types. Furthermore, zUMIs and umis also tended to find more unique DEGs for each cell population than did the other pipelines.

In conclusion, a detailed comparison of the performance of seven high-throughput scRNA-seq data processing pipelines was performed. Overall, different pipelines have their own characteristics in running time performance, computational resource requirements, and data analysis quality. Thus, it is important for researchers to understand these differences to make an informed selection of the proper tools for their scRNA-seq studies.

Our evaluation study has limitations. First, since the ground truthing of the real data was usually unknown or not reliable, biological analysis in many cases was performed based on comparisons among different pipelines. For this reason, some of the results in this study are not conclusive. Second, the biological analysis comparisons performed in this study were by no means comprehensive. Many other aspects of scRNA-seq studies, such as highly variable gene identification, trajectory inference, and gene-gene interactions, were not included in this study due to limitations in scope. Nevertheless, we made all seven pipelines and the evaluation scripts used in this study, which have been encapsulated within Nextflow and Docker for easy deployment, available on-line at https://github.com/xmuyulab/scRNAseq_pipelines so that users can deploy these tools for their own data analysis tasks and perform similar comparisons on their own data to gain better insight into the performances of the different pipelines.

