## Supplementary for "Comparison of High-Throughput Single-Cell RNA Sequencing Data Processing Pipelines"

**Table S1.** Characteristic summary of the high-throughput scRNA-seq data processing pipelines investigated.

| | Drop-seq-tools (2015) | UMI-tools (2017) | umis (2017) | Cell Ranger (2017) | zUMIs (2018) | dropEst (2018) | scPipe (2018) |
|---|---|---|---|---|---|---|---|
| **scRNA-seq Platforms*** | UMI-based protocols | UMI-based protocols | almost all | 10X Genomics | almost all | droplet-based protocols | UMI-based protocols |
| **Open Source** | no | yes | yes | yes | yes | yes | yes |
| **Programming Language** | Java | Python | Python | Python, Rust | Perl, R | C++, R | C++, R |
| **Transcript Sequence QC** | yes | no | no | yes | no | no | no |
| **Cell Barcode Correction** | no | no (default) /Hamming-distance | no | Hamming-distance | no | Hamming-distance | no |
| **UMI Correction** | Hamming-distance | graph-based method | none | base quality & Hamming-distance | Hamming-distance | Bayesian model | Hamming-distance |
| **Aligner** | STAR | STAR+^ | Kallisto, Rapmap | STAR | STAR+ | STAR+ | Subread+ |
| **Statistics Report** | text summary | - | - | web report & downstream analysis | - | - | web report |
| **Intron Mapping** | no | no | no | no | yes | yes | no |
| **Cell Barcode Number Determination** | manual (cell number / white list) | manual (cell number / white list) | manual (cell number / white list) | manual (cell number) | automatic/ manual (cell number / white list) | automatic | manual (low limit of reads / cell number) |
| **Cited#** | 1511 | 111 | 136 | 560 | 20 | 11 | 11 |

* *Droplet-based protocols include 10X, CEL-Seq2, Drop-seq, iCLIP, inDrop, Seq-Well, SPLiT-seq. UMI-based protocols further extend the range of droplet-based protocols to include any protocols in which UMI is involved in sequencing library construction. ZUMIs and umis are compatible with almost all sequencing platforms with or without UMI.*

^ *"+" indicates that the pipeline also supports other aligners in additional to the mentioned one.*

### *Cited data were collected on Web of Science in December 2019. Only the papers in which the corresponding pipelines were introduced for the first time are considered here.*

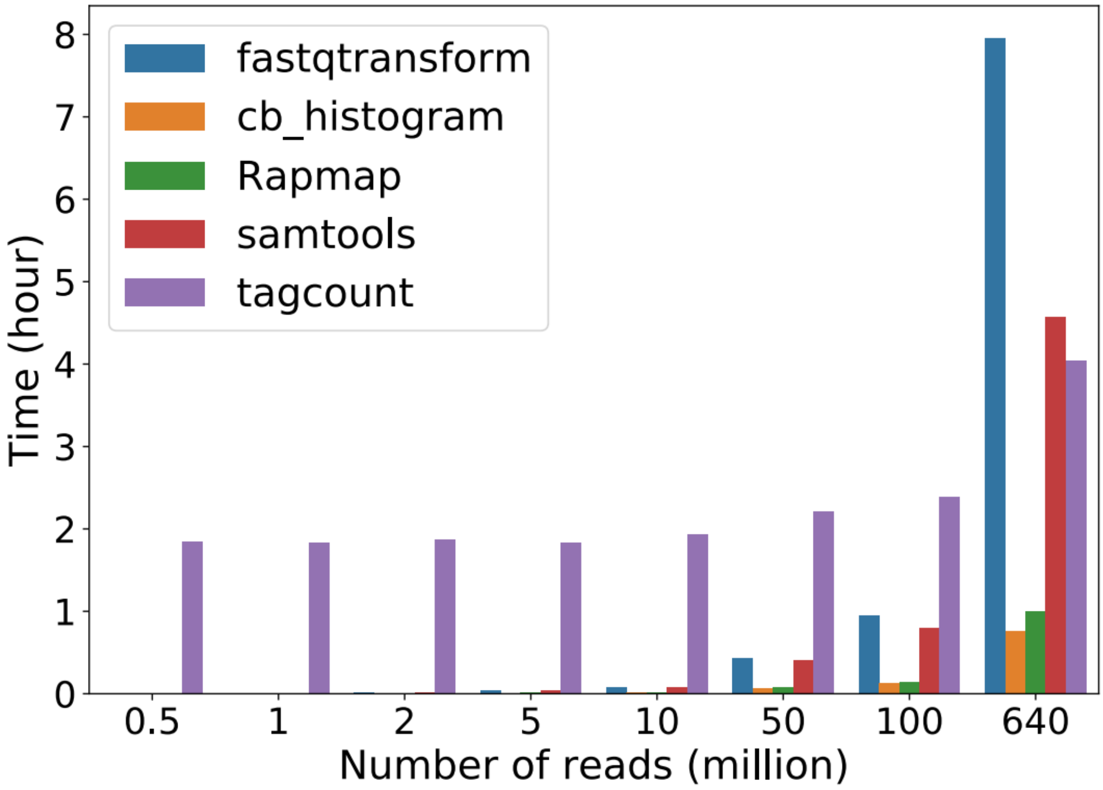


**Figure S1.** Running time of individual steps of *umis* on 10X-PBMC-10k dataset.

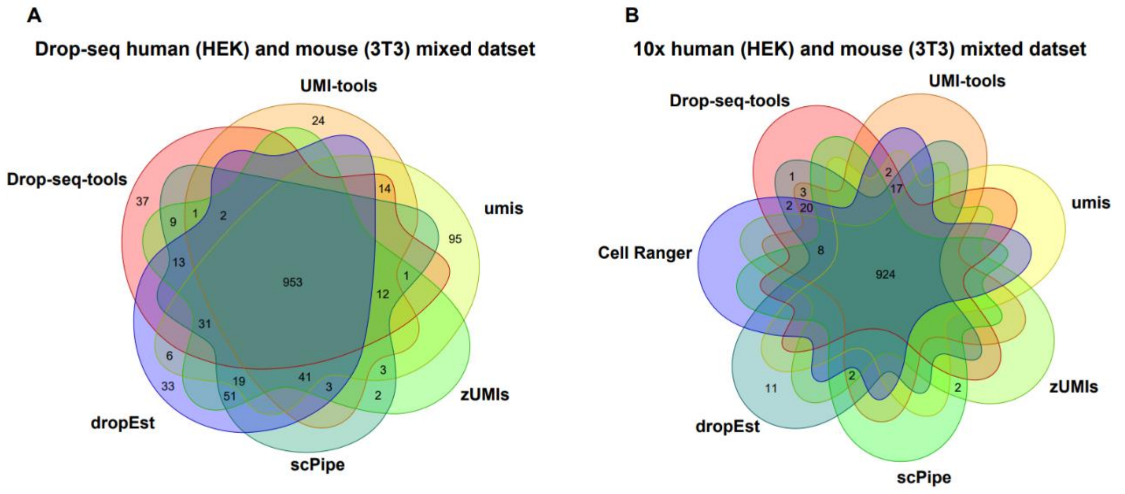


**Figure S2.** Venn diagrams of cell barcodes generated by different pipelines. (**A**) Drop-HM dataset. (**B**) 10X-HM dataset.

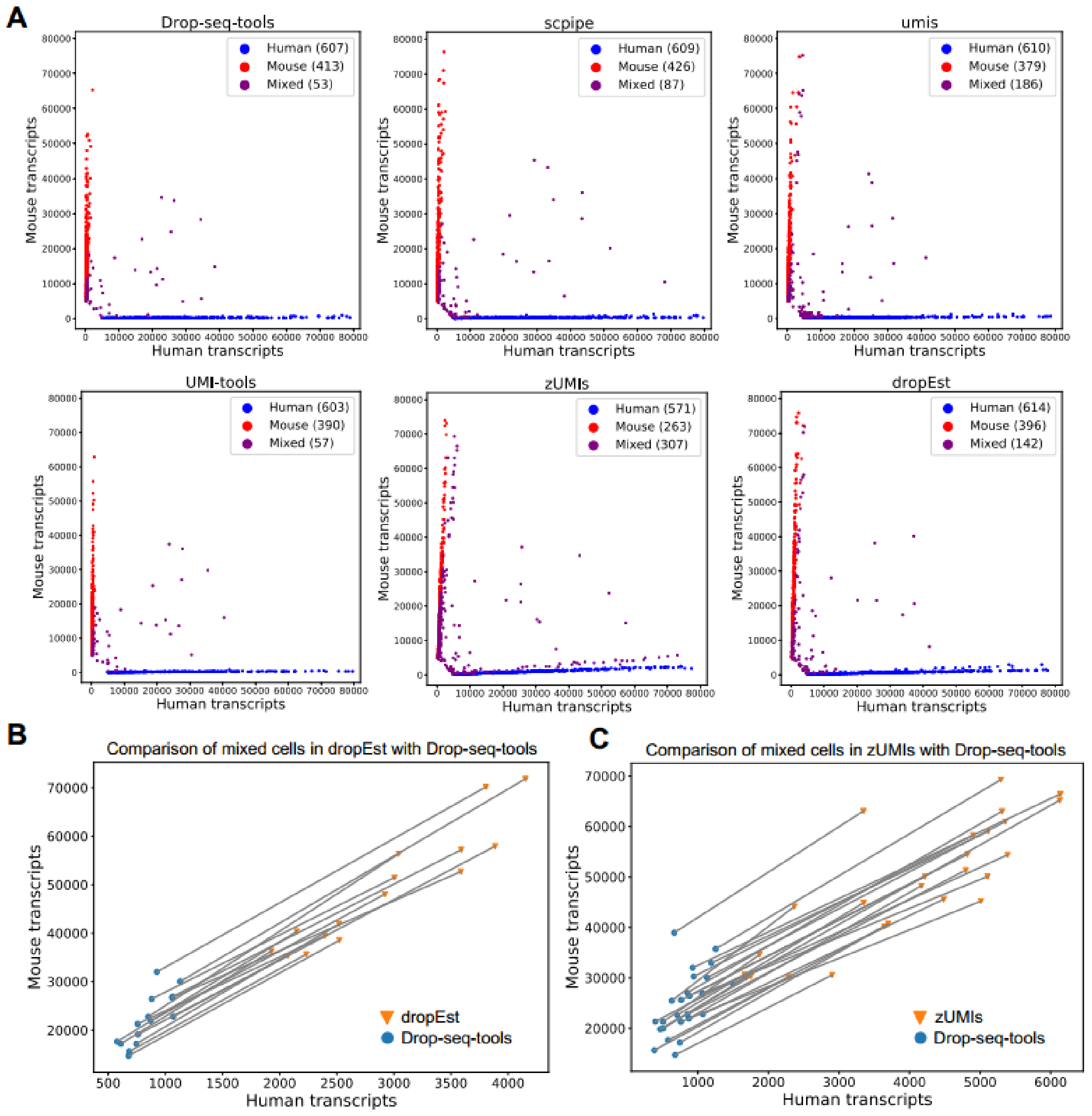


**Figure S3.** Transcript quantification analysis of Drop-HM dataset. (A) Scatter plots of the number of human and mouse transcripts of each cell barcode. (B) and (C) Difference of the number of human and mouse transcripts of (B) dropEst and (C) zUMIs compared with Drop-seq-tools. A pair of points linked by a single line stands for the same cell processed by two different pipelines. Points shown in the figures are specified subsets of mixed cells in corresponding pipelines.

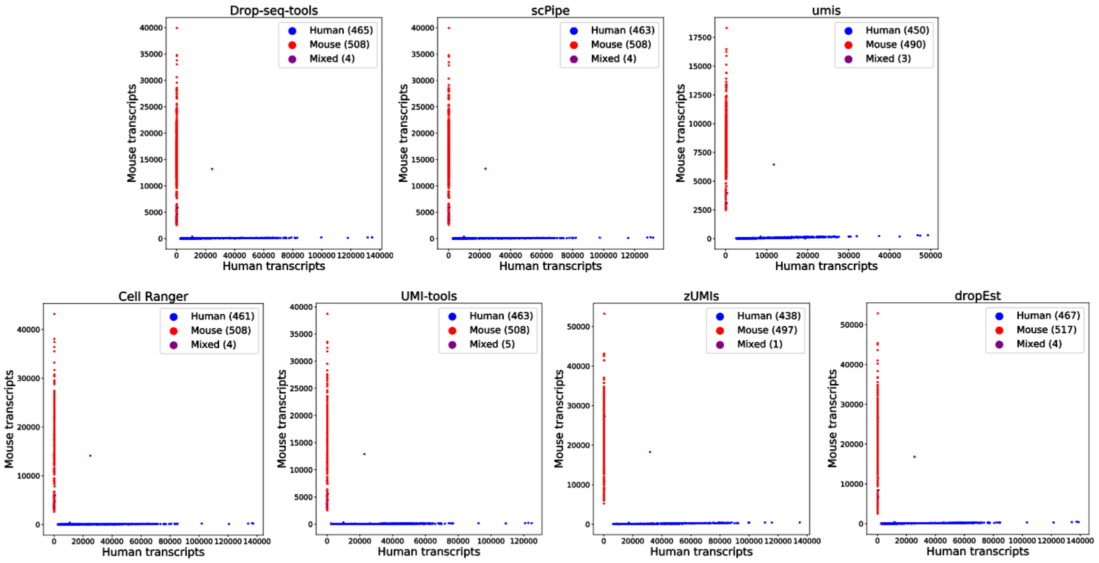


**Figure S4.** Scatter plots of the number of human and mouse transcripts of each cell barcode on 10X-HM dataset.

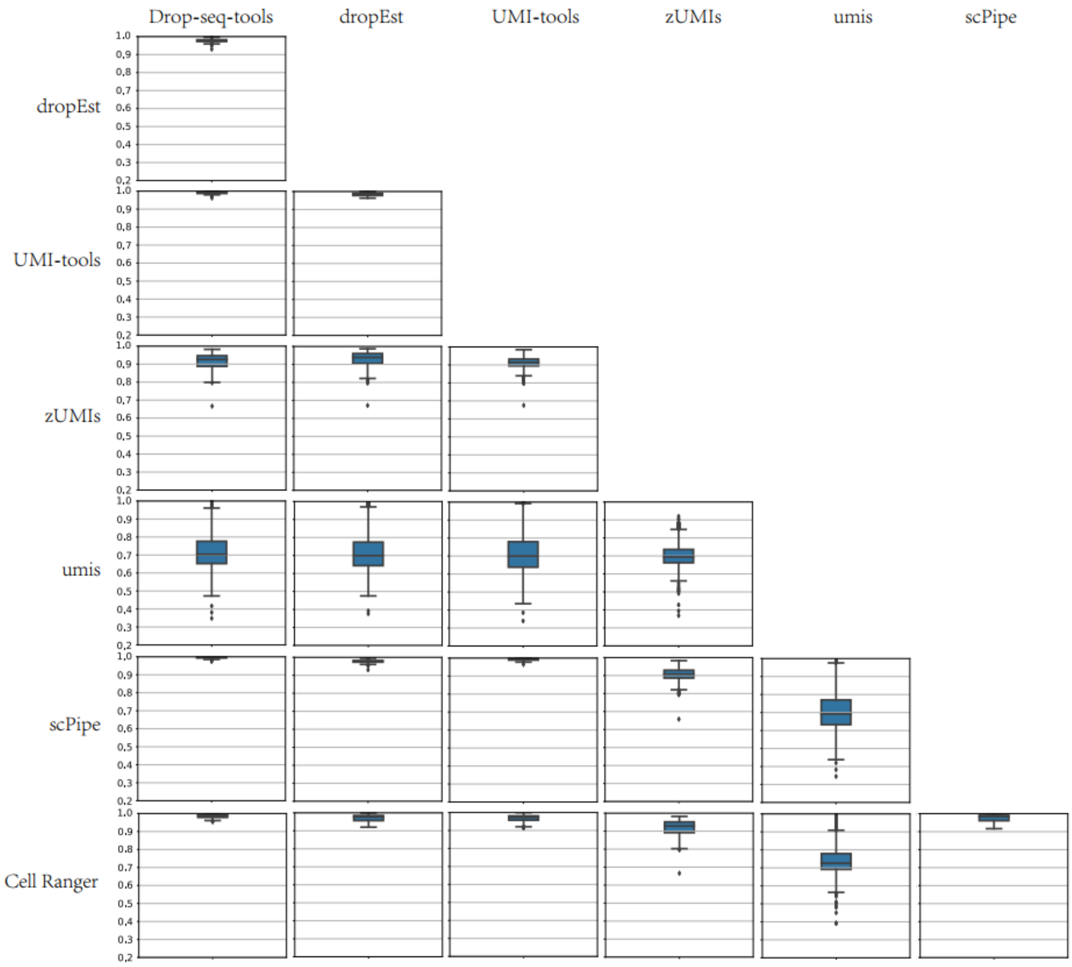


**Figure S5.** Pairwise Pearson boxplots of the gene expression of 924 shared cells across seven pipelines on 10X-HM dataset.

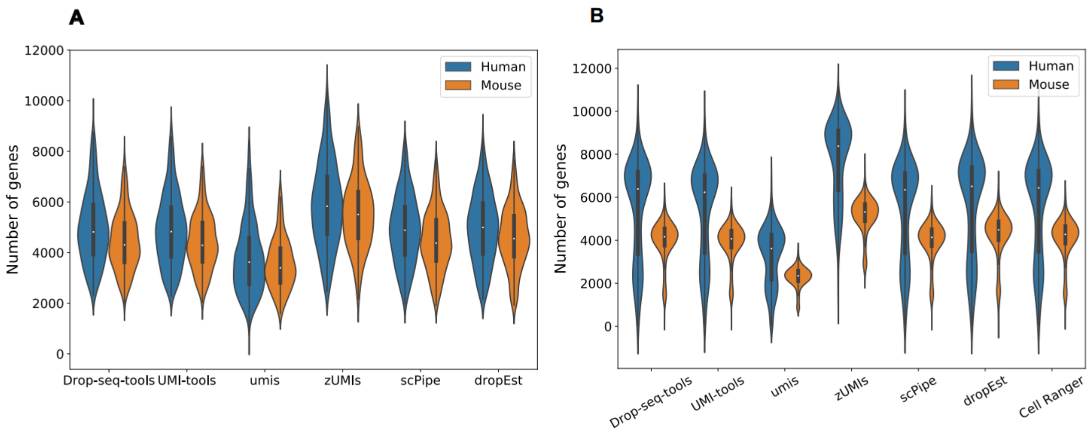


**Figure S6.** Comparison of number of genes detected by different pipelines on (**A**) Drop-HM dataset, (**B**) 10X-HM dataset.

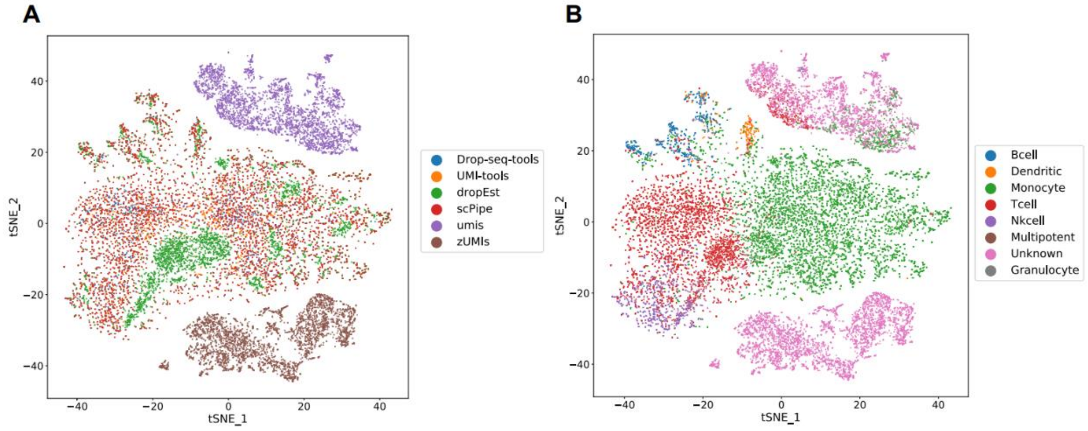


**Figure S7.** TSNE plots of the same cells of Seq-Well-PBMC dataset generated by different pipelines. (**A**) coloured by pipelines. (**B**) coloured by cell types identified with *SuperCT* web application. *Cell Ranger* was not tested due to compatibility issues.

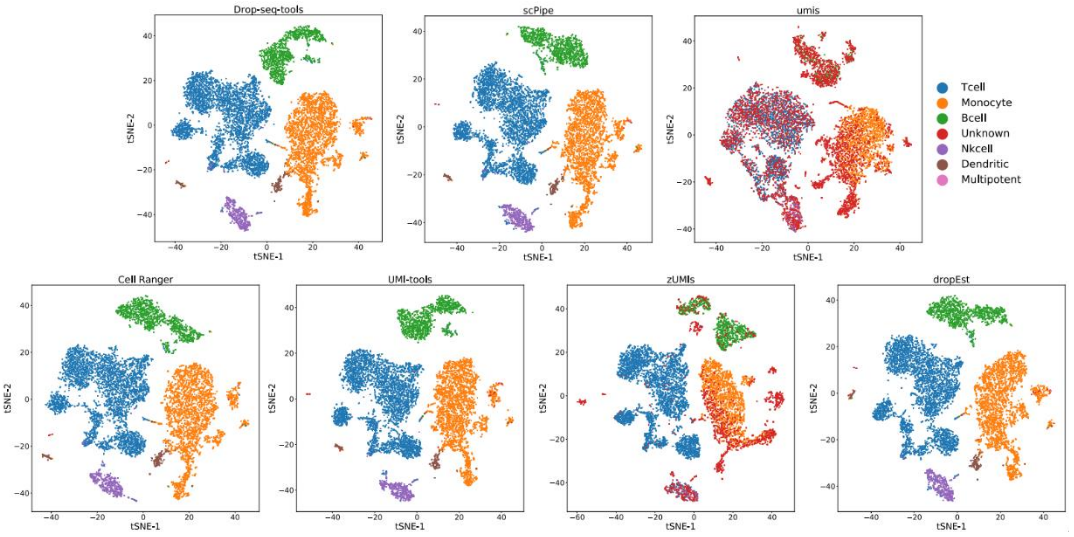


**Figure S8.** TSNE plots of 10X-PBMC dataset by *Seurat* based on the expression matrices generated by different pipelines. The cell types were identified with *SuperCT* web application.

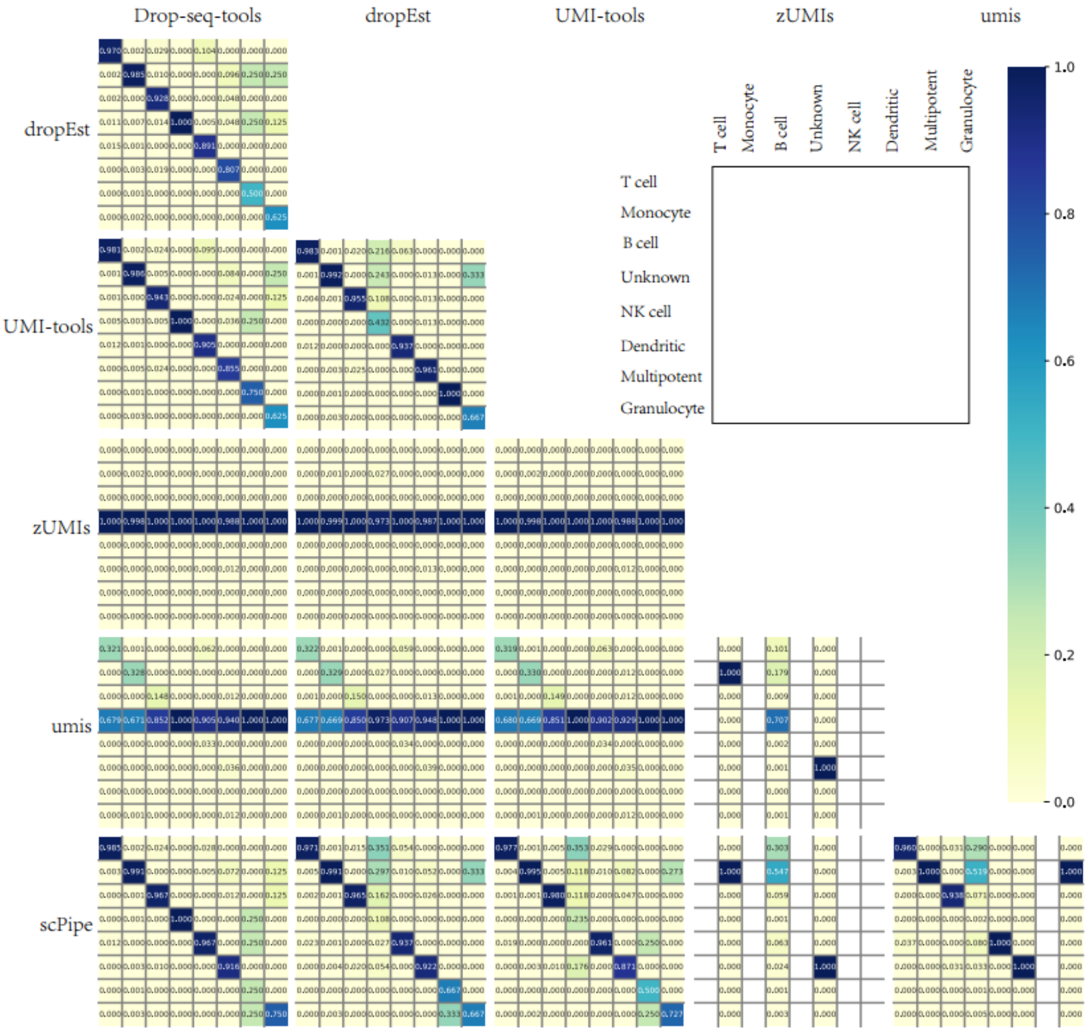


**Figure S9.** Confusion matrices of the cell types of Seq-Well-PBMC dataset identified with *SuperCT* web application across six pipelines.

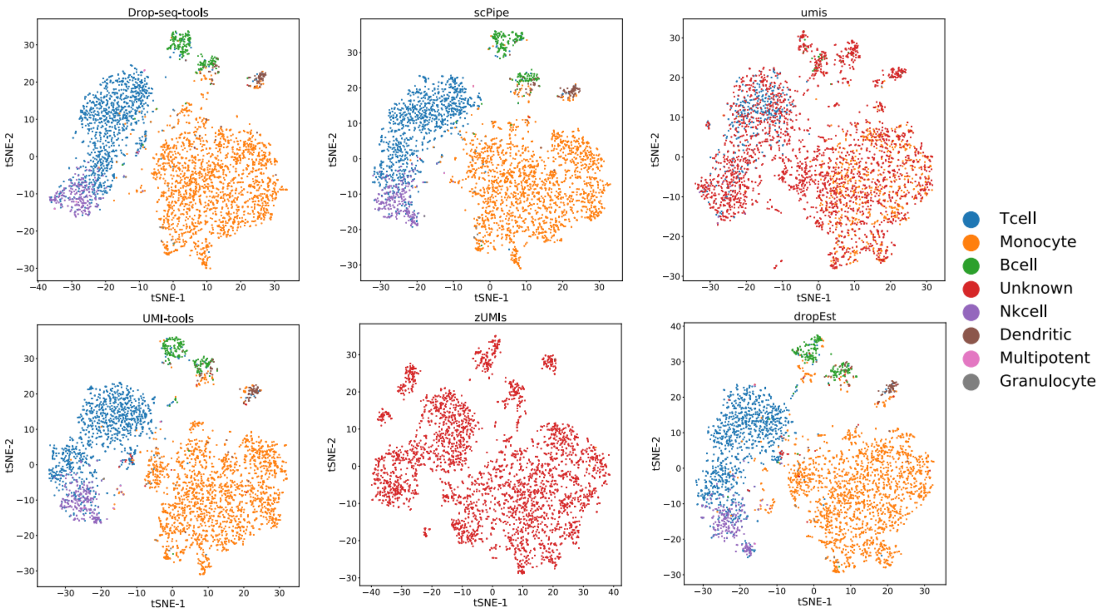


**Figure S10.** TSNE plots of Seq-Well-PBMC dataset by *Seurat* based on expression matrices generated by different pipelines. The cell types were identified with *SuperCT* web application.

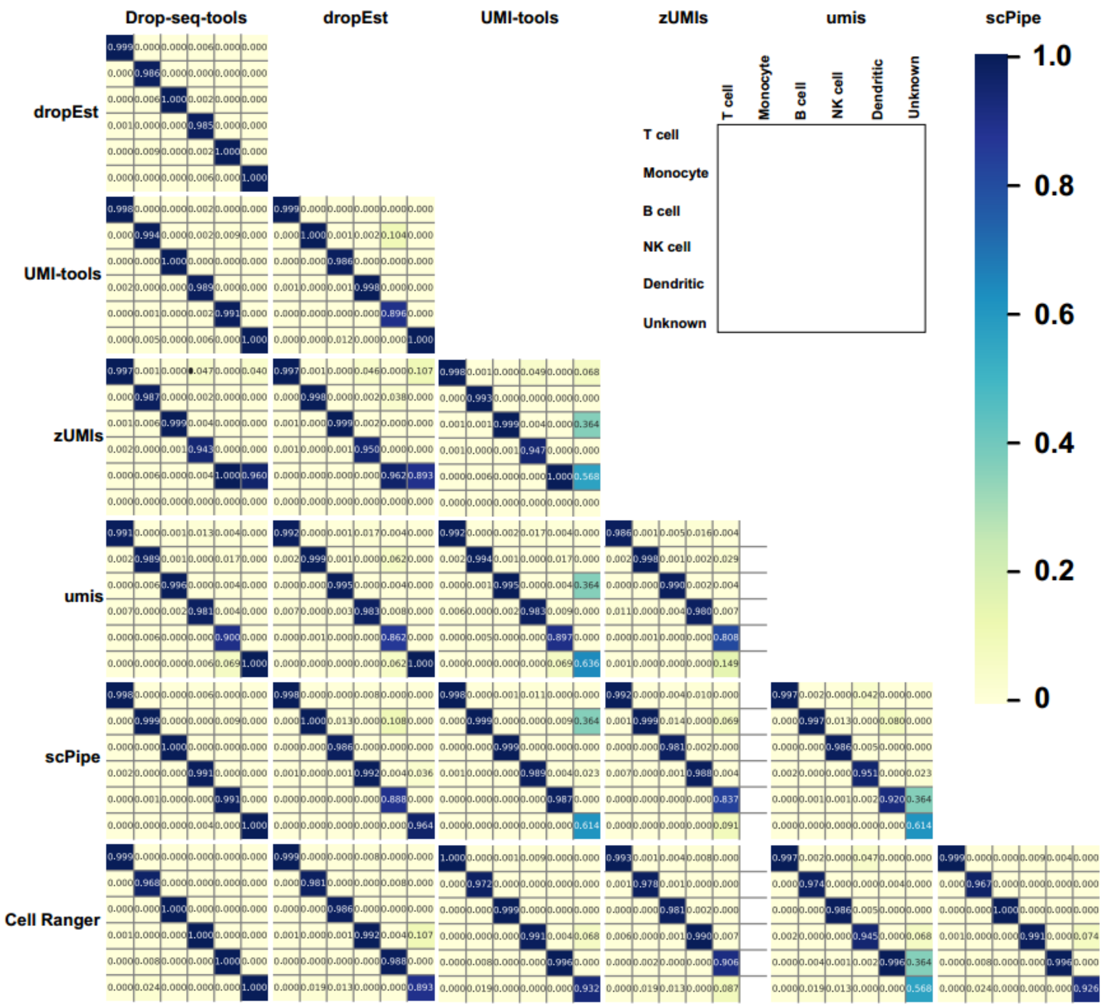


**Figure S11.** Confusion matrices of the cell types of 10X-PBMC-10k dataset identified by majority voting of unsupervised clustering-based methods by three independent researchers.

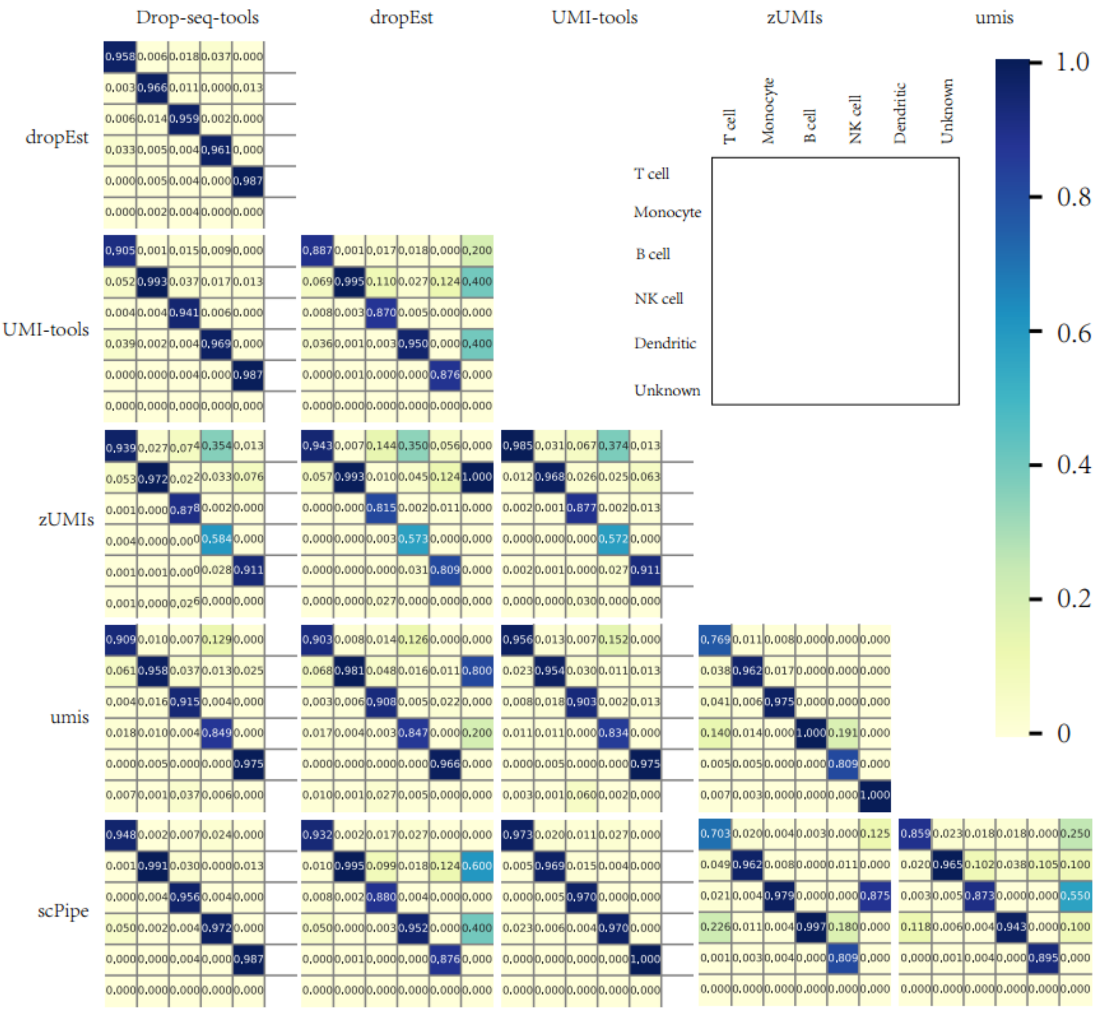


**Figure S12.** Confusion matrices of the cell types of Seq-Well-PBMC dataset identified by majority voting of unsupervised clustering-based methods by three independent researchers.

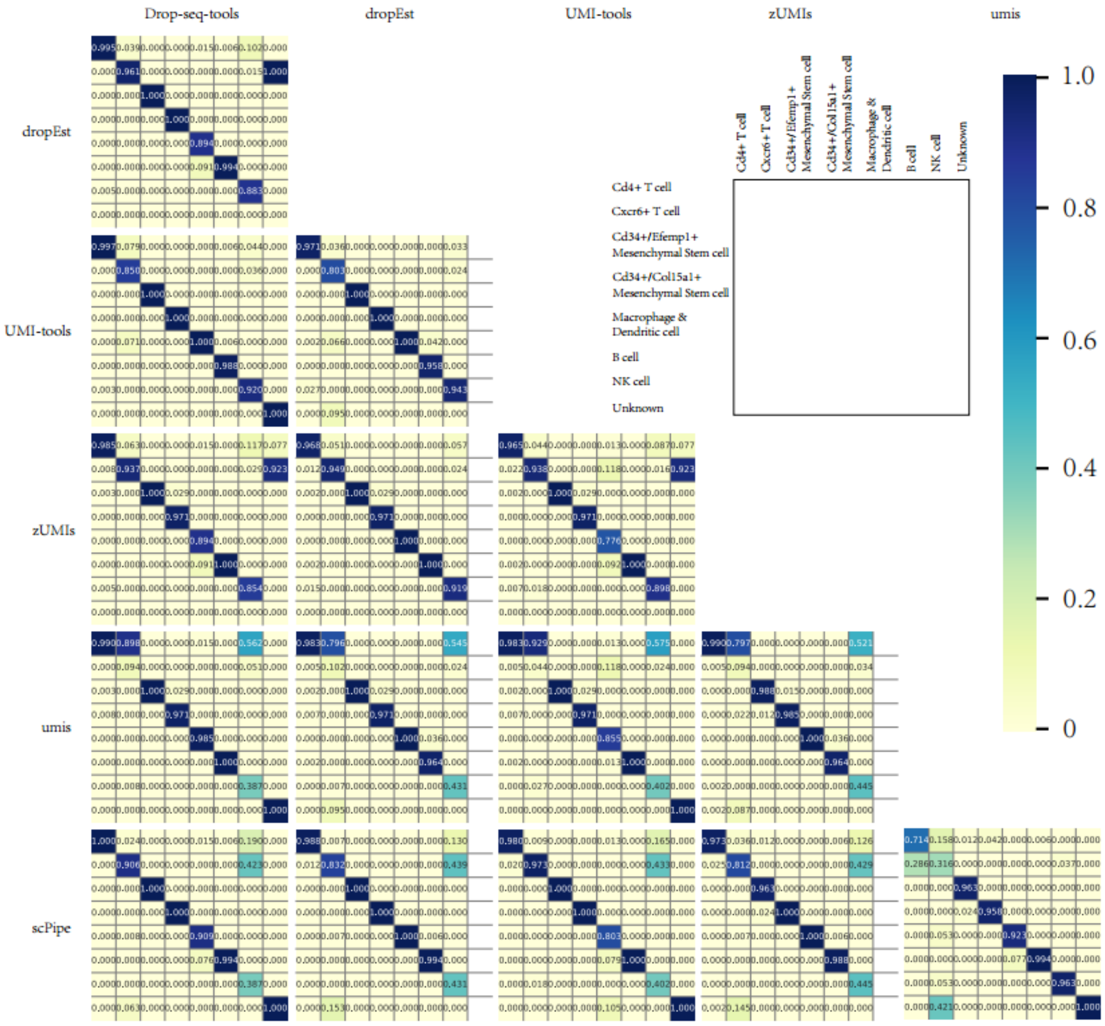


**Figure S13.** Confusion matrices of the cell types of Quartz-SVF dataset identified by majority voting of unsupervised clustering-based methods by three independent researchers.

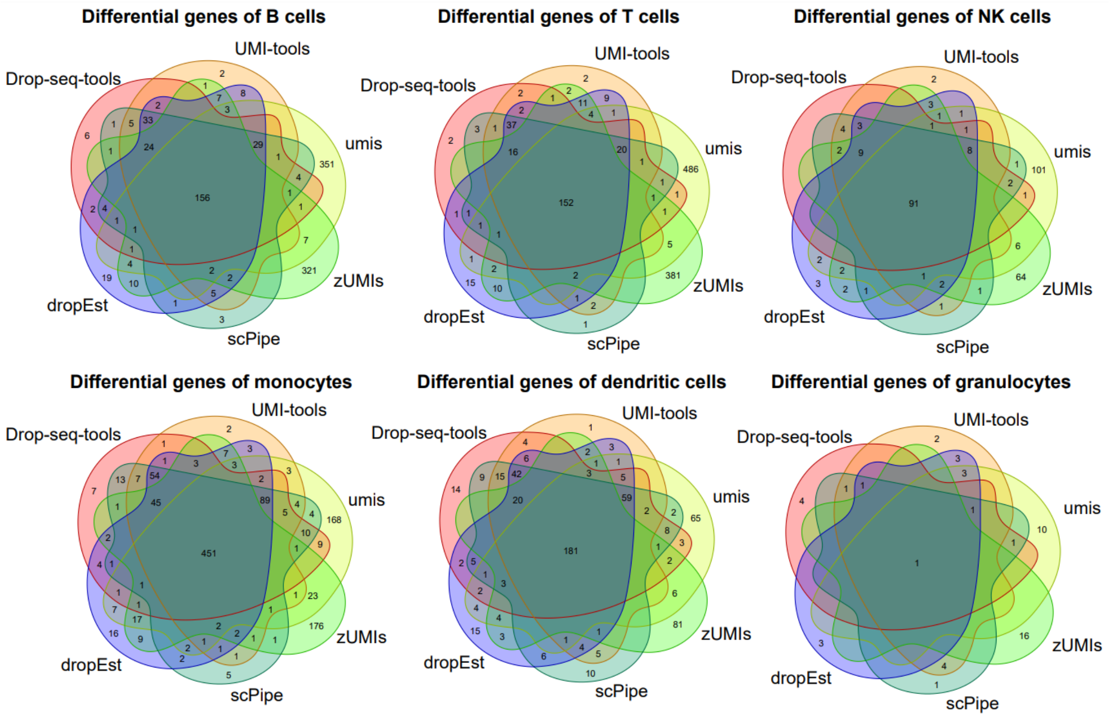


**Figure S14.** Venn diagrams of differential expressed genes found by *Seurat* with adjusted p-value (Bonferroni correction) less than 0.05 in Seq-Well-PBMC dataset.
